# The *rx3* gene contributes to the evolution of eye loss in the cavefish *Astyanax mexicanus*

**DOI:** 10.1101/2024.12.11.628050

**Authors:** Devin Shennard, Itzel Sifuentes-Romero, Rianna Ambosie, Jennah Abdelaziz, Erik R. Duboue, Johanna E. Kowalko

## Abstract

Uncovering mechanisms by which sensory systems evolve is critical for understanding how organisms adapt to a novel environment. *Astyanax mexicanus* is a species of fish with populations of surface fish that inhabit rivers and streams and cavefish that have adapted to life within caves. Cavefish have evolved sensory system changes relative to their surface fish counterparts, providing an opportunity to investigate mechanisms underlying sensory system evolution. Here, we report the role of the gene *retinal homeobox 3* (*rx3*) in cavefish eye evolution. We generated surface fish with putative loss-of-function mutations in the *rx3* gene using CRISPR-Cas9 to determine the role of this gene in eye development in this species. These *rx3* mutant surface fish fail to develop eyes, demonstrating that *rx3* is required for surface fish eye development. Further, *rx3* mutant surface fish exhibit altered behaviors relative to wild-type surface fish, suggesting that the loss of eyes impacts sensory-dependent behaviors. Finally, eye development is altered in cave-surface hybrid fish that inherit the mutant allele of *rx3* from surface fish relative to siblings that inherit a wild-type surface fish *rx3* allele, suggesting that cis-regulatory variation at the *rx3* locus contributes to eye size evolution in cavefish. Together, these findings demonstrate that, as in other species, *rx3* is required for eye development in *A. mexicanus*. Moreover, they suggest that variation at the *rx3* locus plays a role in the evolved reduction of eye size in cavefish, shedding light on the genetic mechanisms underlying sensory system evolution in response to extreme environmental changes.

## 1. Introduction

Animals that inhabit different environments often have access to drastically different sensory cues. Evolution of sensory systems has led to organisms being able to distinguish ecologically relevant cues in their environments, and has resulted in sensory system diversity across species (Oteiza & Baldwin, 2021). For example, compared to visually-foraging chickens, ducks which forage using tactile cues have a larger proportion of mechanoreceptive neurons in the trigeminal ganglia, which are responsible for sensing mechanical stimuli in the bill (E. R. Schneider et al., 2017). Further, tactile foraging birds have an expansion of the size of the principal trigeminal nucleus, the part of the brain that processes input from tactile information during feeding, relative to visually foraging birds (Gutiérrez-Ibáñez et al., 2009). Similarly, multiple aspects of the visual system differ between nocturnal and diurnal birds of prey, which are thought to contribute to differences in visual acuity in low-light versus daylight (Potier et al., 2020). These studies reveal how ecological challenges can lead to evolution of sensory systems, however, the genetic and developmental mechanisms driving these adaptations remain largely unresolved. Identifying the mechanisms by which sensory systems evolve is critical to understanding how organisms adapt to novel environments.

*Astyanax mexicanus* is a freshwater fish species that is a powerful model system for investigating mechanisms contributing to trait evolution. *A. mexicanus* consists of two morphs, a river-dwelling surface fish and a cave-dwelling fish. At least two independent lineages of sighted surface-dwelling fish have repeatedly colonized caves in Mexico, giving rise to over 30 cavefish populations that have evolved a number of adaptations that allow them to live their dark cave environments (Bradic et al., 2012; Garduño-Sánchez et al., 2023; Herman et al., 2018; Moran et al., 2023). These adaptations include changes to cavefish sensory systems, including regression of the eyes, expansion of the lateral line, enhancement of olfaction, and an increase in the number and distribution of tastebuds (Bensouilah & Denizot, 1991; Berning et al., 2024; Blin et al., 2018; Franz-Odendaal & Hall, 2006; W. R. Jeffery, 2005; Lunsford et al., 2022; Shiriagin & Korsching, 2019; Sifuentes-Romero et al., 2023; Teyke, 1990; Varatharasan et al., 2009). The phenotypic divergence in sensory systems in a single species make *A. mexicanus* a powerful model for studying sensory system evolution.

Among evolved sensory systems in cavefish, eye regression has been studied extensively. Cavefish initiate eye development, however, early eye development in cavefish is distinct from that of surface fish in a number of ways, including reductions in the size of the developing optic cup and lens relative to surface fish, as well as loss of the ventral retina (William R. Jeffery et al., 2003; Pottin et al., 2011; Yoshiyuki Yamamoto et al., 2004). Shortly after the optic cup is formed in cavefish, eye regression begins with lens apoptosis, which is followed by apoptosis of the retina, and which ultimately leads to degeneration and the eyeless phenotype seen in adulthood (Alunni et al., 2007; William R. Jeffery & Martasian, 1998; Strickler et al., 2007; Y. Yamamoto & Jeffery, 2000). Many of these alterations to development are shared across *A. mexicanus* cavefish populations (William R. Jeffery et al., 2003; Sifuentes-Romero et al., 2020).

In addition to developmental changes that characterize cavefish eye regression, the genetic underpinnings of cavefish eye regression have been investigated, primarily through crosses and genetic mapping studies. Cavefish and surface fish are interfertile and can produce hybrid progeny that have been used for genetic mapping of traits that have evolved in cavefish, including eye loss (K. O’Quin & McGaugh, 2016; Wiese et al., 2024). These mapping studies have revealed that multiple genetic loci contribute to differences in eye size, pupil size, and alterations to the retina in cavefish compared to surface fish (Borowsky, 2016; Kowalko, Rohner, Linden, et al., 2013; Kowalko, Rohner, Rompani, et al., 2013; K. E. O’Quin et al., 2013; Protas et al., 2007, 2008; Warren et al., 2021; Yoshizawa et al., 2012). While sequencing of the *A. mexicanus* genomes has led to the identification of candidate genes underlying eye Quantitative Trait Loci (QTL) in this species (McGaugh et al., 2014; Warren et al., 2021; Wiese et al., 2024), few candidate genes have been functionally interrogated, and the genes and genetic variants that contribute to eye regression in cavefish are largely unknown (for exception, see (Ma et al., 2020). Addressing these gaps is critical to elucidating the genetic and developmental pathways underlying sensory system evolution.

The gene that encodes the transcription factor Retinal homeobox 3 (Rx3), has been identified as a candidate for eye evolution in *A. mexicanus* (McGaugh et al., 2014; Sifuentes-Romero et al., 2020; Warren et al., 2021). The *rx3* gene falls under an QTL for eye size and is differentially expressed between surface and cave fish in early development during specification of the eye field in the anterior neural plate (Agnès et al., 2022; Leclercq et al., 2024; McGaugh et al., 2014; Sifuentes-Romero et al., 2020; Warren et al., 2021). The function of the *rx3* gene (or the homologous RAX gene in mammals) is highly conserved among vertebrates, playing an essential role in eye development from Xenopus to mouse and humans (Bailey et al., 2004; Chuang et al., 1999; Deschet et al., 1999; Mathers et al., 1997; Muranishi et al., 2012). In fish, mutations in *rx3* cause presumptive eye field cells to incorporate into the telencephalon, highlighting the role of *rx3* in cell fate specification at early developmental stages (Stigloher et al., 2006). Further, mutations in *rx3* prevent optic vesicle evagination and consequently result in an eyeless phenotype (Kennedy et al., 2004; Loosli et al., 2003, 2001; Winkler et al., 2000). Together, these findings position *rx3* as a compelling candidate for investigating both the developmental mechanisms of eye formation in surface fish and the evolutionary modifications that contribute to eye reduction in cavefish, and understanding the mechanisms behind the role of this gene in evolution of eyes could have broad impacts to understanding sensory evolution.

Here, we establish CRISPR-generated lines of *rx3* mutant surface fish to investigate the role *rx3* plays in eye development and vision in *A. mexicanus*, as well as the role the cave alleles of *rx3* play in the evolution of eyes in cavefish. Our findings demonstrate that *rx3* is essential for both eye formation and larval behaviors in surface fish. Moreover, we present data that suggests that cis-regulatory variation at the *rx3* locus likely contributes to the evolutionary reduction of eye size in cavefish, highlighting *rx3* as a key player in sensory system evolution.

## 2. Materials and Methods

### 2.1. Fish husbandry and breeding

All *A. mexicanus* fish used in this study were generated from lab-bred stocks that originated in Mexico, and were maintained in the Florida Atlantic University and Lehigh University fish facilities. All fish handling and husbandry was done in accordance with Florida Atlantic University and Lehigh University Institutional Animal Care and Use Committees (IACUC). All juvenile and adult fish were housed at 23°C and kept on a 14:10 light/dark cycle. Larvae were raised at 25°C in glass bowls at densities of approximately 100 fish per bowl up to 6 days post fertilization (dpf) and then transferred to 2-liter tanks at 23°C in densities of 30 fish per tank. Starting at 6 dpf, fish were fed brine shrimp twice a day. Juvenile fish were fed brine shrimp twice a day, as well as gemma (GEMMA Micro 300 from Skretting) once a day. At later juvenile stages, fish were transitioned to being fed gemma and Zeigler pellets (Zeigler Zebrafish Diet SKU # AH271, Pentair), then to pellets only.

### 2.2. Generation of rx3 mutant fish

To generate germline transmitting lines of *rx3* mutant fish, *rx3* crispant surface fish, surface fish that as single cell embryos were injected with a guide RNA (gRNA) targeting the *rx3* gene (gRNA target sequence: 5’-GTGTAGCTGAAACGTGGTGA-3’) and Cas9 mRNA, which were described previously (Warren et al., 2021), were incrossed. At adult stages, offspring from these crosses were fin clipped and DNA was extracted for genotyping, as described previously (Klaassen et al., 2018; Kowalko et al., 2016; Ma et al., 2015). To extract DNA, fin clips were placed into 100 µl of 50 mM NaOH and incubated at 95°C for 30 minutes. Following incubation, 10 µl of 1 M Tris-HCl pH 8 was added to each sample. PCR was performed using the extracted DNA and primers designed to amplify the region of DNA surrounding the gRNA site located in exon 2 of the *rx3* locus (F: 5’-AGCCCGGACCGTAAGAAG-3’ and R:5’-GCTGTAAACGTCGGGGTAGT3’). Heterozygotes were identified via gel electrophoresis following PCR amplification. Next, heterozygotes were confirmed by sequencing the TA cloned PCR amplicon to identify two G1 fish heterozygous for a 10 base pair (bp) deletion in exon 2 of the *rx3* gene. The *rx3* heterozygous G1 fish were crossed with wild-type surface fish and G2 offspring were grown to adulthood and genotyped by PCR and gel electrophoresis following DNA extraction from fin clips. Heterozygous G2 offspring were then incrossed to produce the G3 homozygous mutant, heterozygous, and wildtype fish which were used for all assays here. Surface-Pachón hybrid fish were generated by crossing a surface fish that is heterozygous at the *rx3* locus for the 10 bp deletion to a Pachón cavefish in order to generate surface-Pachón hybrid progeny.

### 2.3. Imaging

Eye morphology was assessed in *rx3* mutant, wild-type and heterozygous surface fish larvae at multiple time points, including during optic vesicle evagination (20-22 somite stage (ss)), 36 hours post fertilization (hpf), 5 dpf, and in adulthood (∼1.5 years old). Larval fish less than 24 hpf were chemically dechorionated (0.1 mg/mL pronase, Sigma) prior to imaging. All embryonic and larval fish were euthanized in MS-222 immediately before imaging. Embryonic and larval fish were mounted and imaged in 2% methyl cellulose. Following imaging, whole embryonic fish and larval fish and fin clips from adult fish were immediately collected to extract genomic DNA for genotyping (see above). Incrossed fish, as well as surface-Pachón hybrid fish, were imaged at 5 dpf and genotyped to measure and compare eye diameter between varying genotypes. All fish were imaged on a Zeiss Stemi 508 dissecting microscope using a Canon Rebel DSLR camera. Eye diameter was measured using FIJI (C. A. Schneider et al., 2012-7). In incrossed surface fish, body length was correlated to eye diameter and so we corrected for effects of body length on eye size by dividing eye diameter by standard length (SL) for comparisons. In surface-Pachón hybrid fish, body length was not correlated to eye diameter, and so eye diameter was compared between genotypes.

### 2.4. Prey capture assays

Fish larvae were assayed for prey capture behavior at 8 dpf using previously described protocols (Lloyd et al., 2018). Briefly, fish were assayed prior to daily feeding. Fish were acclimated to the assay room for at least an hour prior to assay start. Fish were placed in a circular well (15.54 mm diameter) filled with fish water and 10-15 brine shrimp that were prepared 24 hours prior to assay start. Feeding behavior was observed for two minutes by filming from above using a FLIR Grasshopper3 High Performance USB 3.0 Monochrome Camera (Edmund Optics Cat. No. 88-607) with a 12mm HP series lens, 1/1” (Edmund Optics, Cat. No. 86570) and recorded using the Spinview program from FLIR’s Spinnaker SDK. Videos were recorded at 50 frames per second at a resolution of 992 x 1000. All videos were analyzed for prey capture behavior using FIJI (Schindelin et al., 2012). Attempts to catch prey, strikes, were analyzed and categorized as successful, unsuccessful, and unmeasurable. We assessed prey capture behavior as described previously (Lloyd et al., 2018). Total number of strikes, as well as success rate were calculated for each fish. Successful strikes were defined as strike attempts where the fish successfully captured prey, and these strikes were further analyzed to quantify the strike angle and strike distance. Any fish with fewer than 3 successful strikes was excluded from analysis. Strike angle and distance were not recorded for unsuccessful strikes, strike attempts that did not result in the capture of the brine shrimp, or multi-strikes, strikes where larvae struck multiple times in quick succession without recovering in between each strike. However, these strikes were included for the calculations of total strikes and success rate.

Strike angle and distance were measured for successful strikes as described previously (Choy et al., 2024; Lloyd et al., 2018). For strike angle, the frame before the start of the strike attempt was used to measure the angle between the midline of the larva from the base of its head and the center of the prey. Strike distance was measured for the shortest distance between the larva’s head and the brine shrimp. Following the assay, fish were euthanized, DNA was extracted, and genotyping was performed as described above.

### 2.5. Optomotor response assays

Fish larvae were assayed for Optomotor response (OMR) at 8 dpf. All fish were fed and transported into the assay room for acclimation at least an hour prior to assay start. To assay OMR, fish were placed in fish water in a 4 welled plate (Nunc rectangular dishes, Thomas Scientific, item number 1228D90) on a Samsung Tab Active Pro tablet (model number SM-T540) and were acclimated to a white background for 1 minute prior to being exposed to a video with 30 second white background followed by moving black and white lines that are 2 centimeters in width and move at a speed of 1 centimeter per second. Moving lines switched the direction in which they moved every 30 seconds for 5 total switches. The assay was filmed using a FLIR Grasshopper3 High Performance 329 USB 3.0 Monochrome Camera (Edmund Optics Cat. No. 88-607) with a 12mm HP series lens, 330 1/1” (Edmund Optics, Cat. No. 86570) and recorded at 30 frames per second at a resolution of 800×1200 using the Spinview program from FLIR’s Spinnaker SDK.

The distance that each fish moved in the direction of the moving lines was measured across all 5 direction changes as previously described in (Choy et al., 2024). Distance was measured by recording the x coordinate of the fish after each line direction change using the program FIJI (C. A. Schneider et al., 2012-7). The x values immediately before and after each line switch were subtracted from one another to determine the distance the fish swam relative to the direction of the lines. The distances traveled in the direction of the lines after all line direction changes were averaged to produce an OMR index score for each fish, which was calculated by dividing the average distance traveled relative to the lines over the course of the trial by the total arena length (total possible distance traveled). Thus, the OMR index ranges from 1 to - 1, with 1 indicating the fish traveled from one end of the arena to the opposite end in the direction of line movement during each line switch over the entire assay and −1 indicating that fish traveled to the end of the arena opposite of the moving lines after each line switch during the entire assay. Following the assay, fish were euthanized, DNA was extracted and genotyping was performed as described above.

### 2.6. In situ hybridization

The *rx3* antisense probe for in situ hybridization was generated from a previous cloned portion of cDNA previous used to generate an *rx3* in situ probe (Sifuentes-Romero et al., 2020). Following sequence verification, the riboprobe was generated from PCR products using in vitro transcription with SP6 RNA Polymerase and labeled with digoxigenin (DIG) using a DIG labeling mix (Roche). In situ hybridization was performed as described in (Laboissonniere et al., 2018; Sifuentes-Romero et al., 2020). Surface fish, as well as Pachón, Molino, and Tinaja cavefish, were bred in the laboratory and embryos were raised to 36 hpf and 60 hpf. Fish were euthanized by rapid chilling, and fixed with 4% paraformaldehyde for 4 hours at room temperature or overnight at 4°C. Fixed larvae were dehydrated into methanol and stored in 100% methanol at - 20°C. Post rehydration, larvae were bleached, washed in PBS with 0.1% Tween-20 (PBT), and then permeabilized in a 1:1000 dilution of 10 µg/mL proteinase K, washed in PBT. Following fixation in 4% paraformaldehyde, fish were washed in PBT and then placed in pre-hybridization buffer (HM= 50% formamide, 5X SSC, 0.1% Tween-20, 0.05 mg/mL Heparin, pH adjusted to 6.0) for 5 hours at 70°C followed by hybridization with probe overnight at 70°C in hybridization buffer plus probe. Following probe hybridization, larvae were washed at 70°C (1 x 10’ 75% HM in 2X SSC, 1 x 10’ 50% HM in 2X SSC, 1 x 10’ 25% HM in 2X SSC, 1 x 10’ 2X SSC, 2 x 30’ 0.2X SSC) and at then at room temperature (1 x 10’ 75% 0.2X SSC in PBT, 1 x 10’ 50% 0.2X SSC in PBT, 1 x 10’ 25% 0.2X SSC in PBT, 1x 10’ 100% PBT). Larvae were then placed into an overnight incubation in an anti-DIG antibody coupled to alkaline phosphatase (Roche) in blocking buffer (PBT, 5% HISS) at 4°C. The next day, larvae were washed in PBT and moved into an alkaline tris buffer (0.1M Tris pH 9.5, 0.05M MgCl_2_, 0.1M NaCl, 0.1% Tween-20) and then stained with NBT/BCIP (Fisher) in the dark. Staining was stopped via a stop solution containing PBS and Tween (3 x 15’ of 1X PBS pH 5.5, 1% Tween-20) and larvae were transferred to 100% glycerol for imaging. The 36 hpf fish were imaged on a Zeiss Stemi 508 dissecting microscope using a Canon Rebel DSLR camera while 60 hpf fish were imaged using a Nikon Eclipse E800 compound microscope using a Canon Rebel DSLR.

### 2.7. Statistics

Data were first tested for normal distribution using Shapiro-Wilkes normality tests and equal variances using F-tests. T-tests were performed to compare groups with normal distributions and equal variances. Normally distributed data with non-equal variance, were assessed using Welch’s corrected t-test. Comparisons for data that was not normally distributed were performed using a Mann-Whitney test. All graphs were generated and statistical analyses were performed using Graphpad Prism version 10.0.0, with the exception of Supplemental Figure 2, which was graphed in Microsoft Excel.

## 3. Results

### 3.1 Expression of rx3 is conserved between surface and cavefish following eye morphogenesis

Previous work demonstrated the expression of *rx3* is reduced in the anterior neural plate during eye specification (Agnès et al., 2022; Leclercq et al., 2024; Sifuentes-Romero et al., 2020; Warren et al., 2021), however, whether *rx3* expression is altered between cavefish and surface fish at later stages of eye development is unknown. We assessed *rx3* expression across three cave populations by *in situ* hybridization. Within the first 24 hours of development, the early optic cup and lens are established in both cave and surface fish (Hinaux et al., 2011; William R. Jeffery & Martasian, 1998). Following early development of the optic cup and lens, at 36 hours post fertilization, we found that *rx3* mRNA expression is not detectable in the eye in surface fish or cavefish by in situ hybridization (Fig 1A-D’). However, *rx3* expression was observed in fish from all populations in the ventral brain (Fig 1A-D’). At 60 hours post fertilization, *rx3* expression is found in the ventral brain, as well as in the eyes in both surface fish and fish from multiple cave populations (Fig 1E-H). Together, this suggests that while alterations in *rx3* expression in cavefish occur during eye specification (Agnès et al., 2022; McGaugh et al., 2014; Sifuentes-Romero et al., 2020; Warren et al., 2021), expression patterns of *rx3* are similar between cavefish and surface fish at 36 and 60 hpf of development.

**Figure 1.**
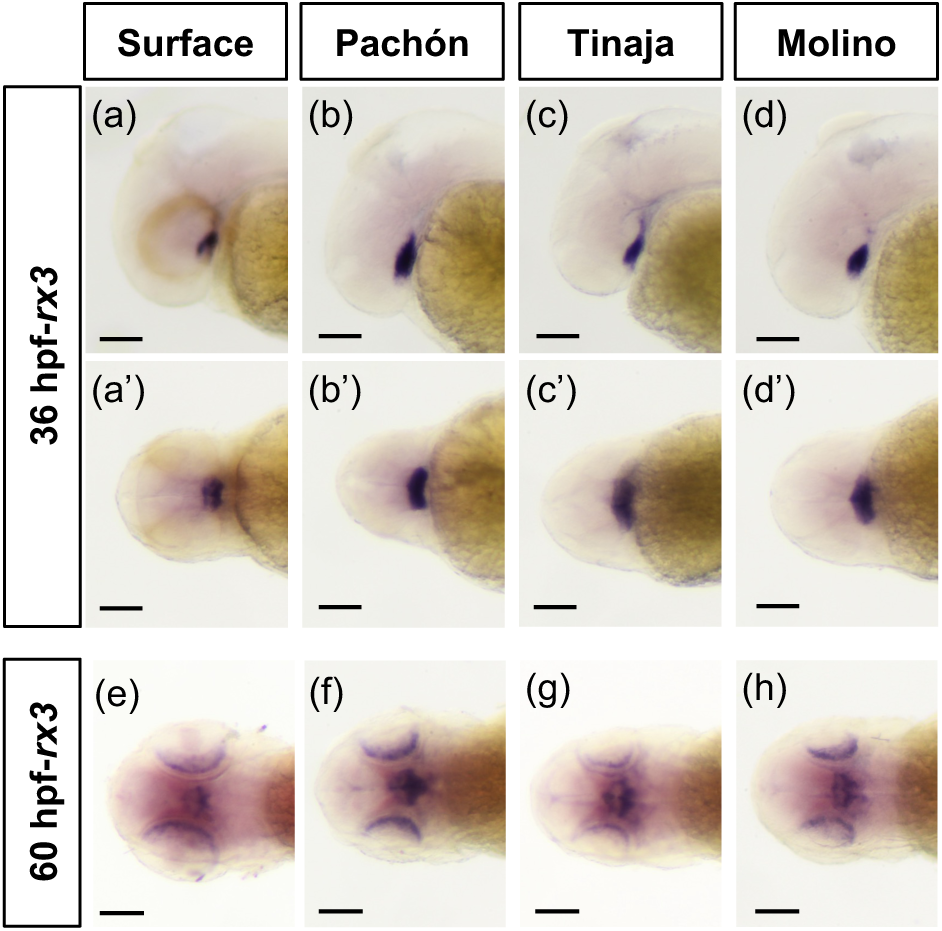
– Characterization of *rx3* expression during early eye development. (a-d’) *rx3 in situ* hybridization at 36 hours post fertilization (hpf) in surface fish (a, a’; lateral and dorsal views, respectively), Pachón (b, b’; lateral and dorsal views, respectively), Tinaja (c, c’; lateral and dorsal views, respectively), and Molino (d, d’; lateral and dorsal views, respectively) cavefish. (e-h) Dorsal views of *rx3 in situ* hybridization at 60 hours post fertilization (hpf) in surface fish (e), Pachón (f), Tinaja (g) and Molino (h) cavefish. All scale bars indicate 0.1 mm.

### 3.2: rx3 mutant surface fish do not develop eyes

The smaller *rx3* expression domain during eye specification in cavefish, combined with the observation that cavefish initially develop smaller eyes than surface fish, together suggest that the reduced *rx3* expression during eye specification plays a role in the reduction of eye size observed in cavefish. To determine the role of *rx3* in eye development in *A. mexicanus*, we first assessed whether *rx3* is required for eye development in surface fish. We generated a line of surface fish heterozygous for a 10-base pair deletion in exon 2 of the *rx3* gene using CRISRP-Cas9. To determine if mutations in *rx3* impact eye development, we incrossed two surface fish heterozygous for the 10-base pair mutation at the *rx3* locus and assessed eye morphology in the offspring.

By 20-somite-stage (20-ss), optic vesicles, which will give rise to both the retina and the retinal pigmented epithelium, are visible in both wild-type cave and surface fish (William R. Jeffery, 2009). While the optic vesicles are visible in wild-type surface fish at 20-22 ss, *rx3* mutant surface fish lack optic vesicles at this stage (Figs 2A&E). By 36 hpf, the optic cup is visible in wild-type surface fish (Fig 2B). In contrast, *rx3* mutant surface fish lack an optic cup at 36 hpf (Fig 2F). This eyeless phenotype in *rx3* mutant surface fish was also observed at later stages in development, at 5 dpf, as well as in adult fish. (Fig 2C, D, G &H). Surface fish heterozygous at the *rx3* locus are eyed, and we found no significant differences in eye diameter size between wild-type surface fish and surface fish heterozygous at the *rx3* locus, suggesting that the eyeless phenotype observed in homozygous mutant surface fish is recessive (Supplemental Fig 1A-B). Together, these data establish *rx3* as a critical gene for early eye development in *A. mexicanus* surface fish and highlight its essential role in initiating the developmental processes required for proper eye formation.

**Figure 2.**
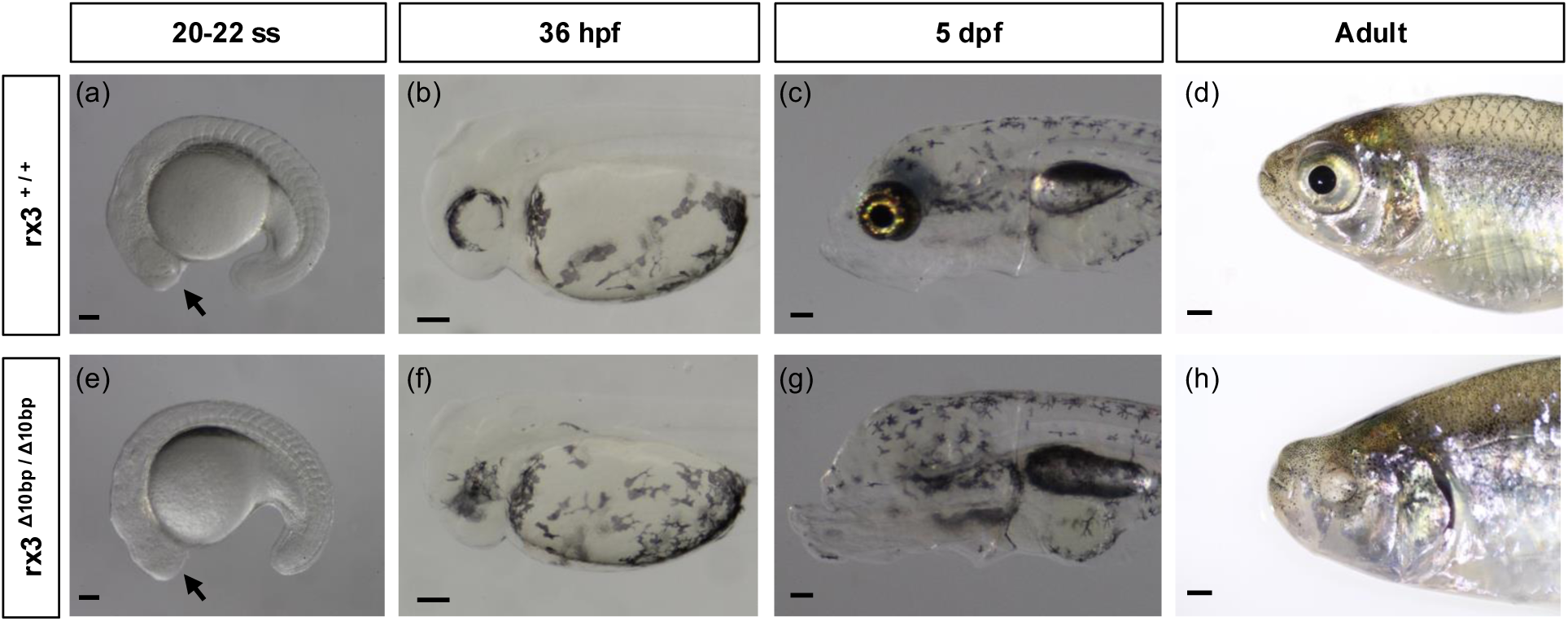
– *rx3* mutant surface fish exhibit an eyeless phenotype. Images of wildtype (*rx3^+/+^*) surface fish (a-d) and sibling *rx3* mutant (*rx3^Δ10bp/Δ10bp^*) surface fish (e-h) at 20-22 somite stage (a,e), 36 hours post fertilization (b,f), 5 days post fertilization (c,g), and adult (d,h) stages. Scale bar is 0.1mm (a-c, e-g) and 1 mm (d,h). Arrow indicates the location of the optic vesicle in wild-type fish and where the optic vesicle would be located in *rx3* mutant fish (a,e).

### 3.3: Multiple behaviors are disrupted in rx3 mutant surface fish

Previous work has characterized differences in multiple sensory-dependent behaviors between surface and cavefish, including behaviors during which surface fish rely on visual cues (Kowalko, Rohner, Rompani, et al., 2013; Lloyd et al., 2018). While loss of vision in cavefish likely plays a role in the evolution of these behaviors, cavefish also exhibit numerous changes in brain anatomy and function that could influence these behaviors (Jaggard et al., 2020; Kozol et al., 2023; Lloyd et al., 2022; Loomis et al., 2019). This raises the question of whether non-visual systems also play a role in driving these behavioral changes. The establishment of a line of *rx3* mutant surface fish that fail to develop eyes provides an opportunity to investigate these behaviors in fish that have lost visual input, allowing us to directly assess the role of eye loss in shaping sensory-dependent behaviors in *Astyanax mexicanus*. Thus, we assessed two visually-mediated behaviors in *rx3* mutant surface fish to determine if surface fish that do not develop eyes behave similarly to cavefish. The optomotor response (OMR) assay quantifies an innate behavior in which fish swim in the direction of moving visual stimuli (Clark, 1981; Neuhauss, 2003; Neuhauss et al., 1999). To determine if OMR is disrupted in *rx3* mutant surface fish, eyeless and eyed surface fish siblings were exposed to moving line stimuli (Fig 3A), and the OMR index was calculated for each fish based on the displacement relative to the direction of line movement (see methods) such that an OMR index score close to 1 indicates that a fish is following the lines, while an OMR index score of close to zero indicates that fish swim without regard to the direction of the traveling lines. Wild-type surface fish larvae had a high OMR index, with the majority of the fish having an OMR index of close to one, suggesting they were able to engage the visually dependent task (Fig 3B). The average OMR index for *rx3* mutant surface fish was significantly lower than that of wild-type surface fish siblings, and was close to zero (Fig 3B), suggesting that *rx3* mutant surface fish have a defective OMR. These findings demonstrate that the loss of eyes in *rx3* mutant surface fish disrupts this visually mediated-behavior and suggests that eye development is essential for engaging in this visual task.

**Figure 3.**
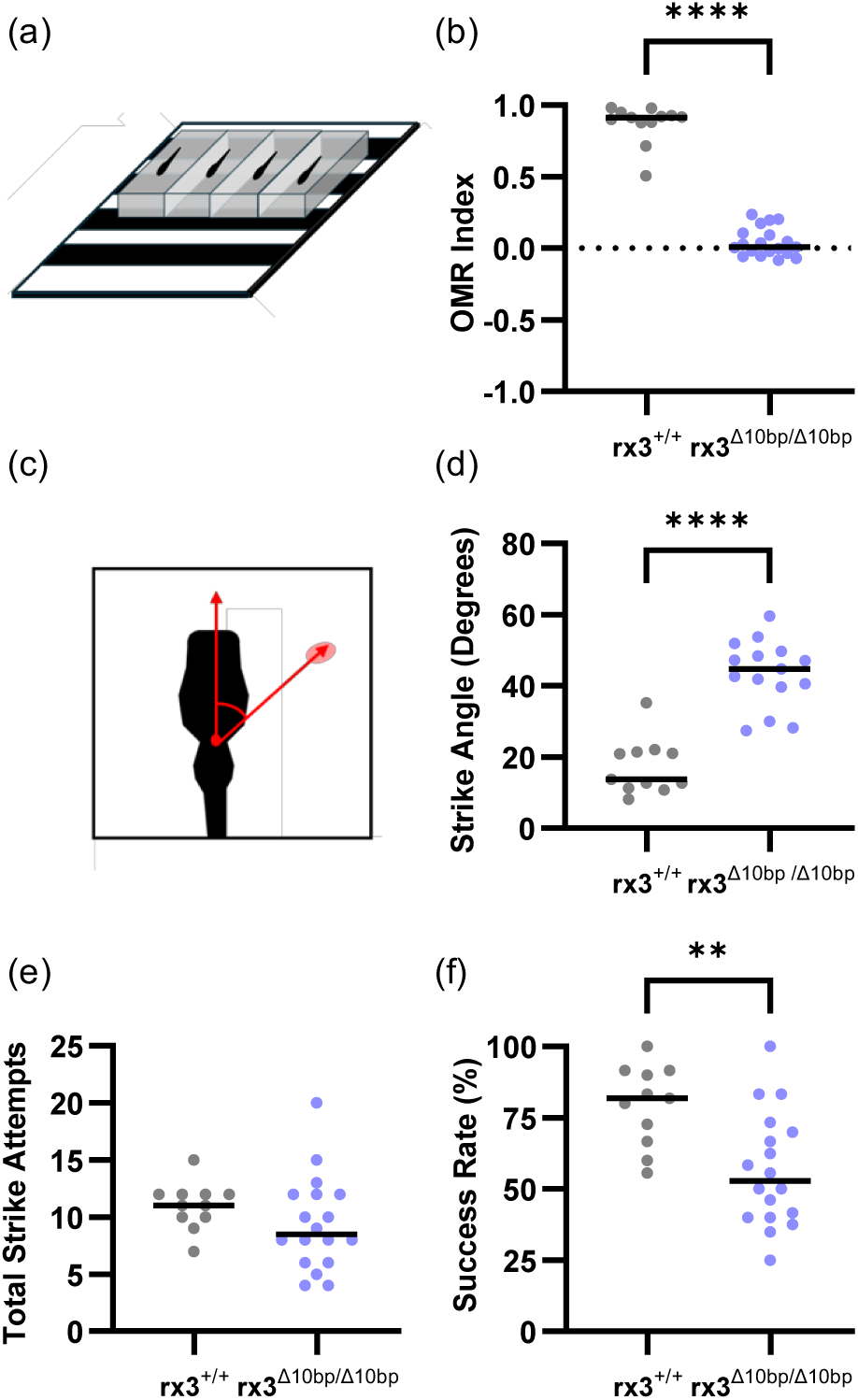
– Visually mediated behaviors are disrupted in *rx3* mutant surface fish. (a) Diagram of optomotor response assay. Fish are placed in wells and exposed to a visual stimulus of moving lines. (b) Optomotor index (OMR Index) is a measure of optomotor response. OMR Index is calculated as the net distance each fish moves between line direction switches in the direction of the lines across multiple line switches such that an OMR index of 1 indicates movement across the whole length of the well in the direction of the lines after every line switch, an OMR index of −1 indicates movement across the whole length of the well in the opposite direction of the lines after every line switch, and an OMR index of 0 indicates movement without regard to the lines or no net movement. OMR index is shown for wild-type (*rx3^+/+^*) and *rx3* mutant (*rx3^Δ10bp/Δ10bp^*) surface fish. Individual points indicate OMR indices for individual fish. Lines indicate the median for each group. OMR index is significantly different for wild-type versus rx3 mutant fish (Mann-Whitney test: n=12,19; P value= <0.0001). (c) Diagram demonstrating how strike angle is measured between fish (black outline) and prey (red oval). (d) Average strike angles of wild-type (*rx3^+/+^*) and *rx3* mutant (*rx3^Δ10bp/Δ10bp^*) surface fish. Individual points indicate average strike angle for individual fish, and line indicates the median per population. Unpaired t-test: n=11,15; t=7.580, p-value<0.0001. (e) Total strike attempts for wild-type (*rx3^+/+^*) and *rx3* mutant (*rx3^Δ10bp/Δ10bp^*) surface fish. Individual points indicate total number of strike attempts for individual fish, and line indicates the median per population. Welch’s t-test: n=11,18; t=1.355, p-value= 0.1870. (f) Success rate, defined as percent of total strikes that are successful, for wild-type (*rx3^+/+^*) and *rx3* mutant (*rx3^Δ10bp/Δ10bp^*) surface fish. Individual points indicate percent of total strikes that were successful for individual fish, and line indicates the median per population. Unpaired t-test: n= 11,18; t=3.318, p-value= 0.0026. ** = p<0.01, **** = p<0.0001.

Surface fish feeding display differences in prey capture behaviors compared to cavefish when visual cues are available. In the light, larval surface fish strike prey head on. In contrast, surface fish in the dark, as well as cavefish, use their lateral line to locate prey and strike from the side (Lloyd et al., 2018). These differences in prey capture can be quantified by measuring strike angle, the angle between the fish and the prey immediately prior to the fish initiating a strike (Lloyd et al., 2018) (Fig 3C). To determine if *rx3* mutant surface fish show altered prey capture dynamics, we quantified prey capture behavior in *rx3* mutant surface fish and wild-type siblings. The *rx3* mutant surface fish struck prey from wide angles, resulting in larger strike angles compared to their wild-type siblings (Fig 3D). While there were no significant differences between the two genotypes in average total number of times fish struck at prey (Fig 3E), the rate of successfully capturing prey was significantly reduced in the *rx3* mutant fish compared to wild-type siblings (Fig 3F), suggesting that visual defects due to eye loss in *rx3* mutant fish impact both prey capture dynamics, and their feeding success. Together, these data demonstrate that multiple behaviors are disrupted in the *rx3* mutant surface fish, suggesting that development of the eyes are required for these behaviors in surface fish.

### 3.4: Eye size in surface-Pachón hybrid fish suggests cis-regulatory changes at the rx3 locus contribute to evolved reductions in eye size in cavefish

The requirement of *rx3* for eye development in surface fish (Fig 2), together with the reduced size of the expression domain of *rx3* in cavefish during eye specification (Agnès et al., 2022; Sifuentes-Romero et al., 2020; Warren et al., 2021), and smaller eye primordia in cavefish during development suggest that the changes in *rx3* expression in cavefish could contribute to the evolution of reduced eye size. One possible explanation for reduced *rx3* expression in the anterior neural plate of cavefish is that factors upstream of *rx3* which regulate the *rx3* expression domain size are different between cave and surface fish. Alternatively, cis-regulatory variation at the *rx3* locus itself could lead to reduced expression in cavefish compared to surface fish. To distinguish between these possibilities, we performed a genetic complementation test by crossing surface fish that are heterozygous at the *rx3* locus for the engineered 10bp *rx3* deletion with Pachón cavefish and quantifying eye size in the resulting cave-surface hybrid progeny. Eye size and morphology varied significantly between individual cave-surface hybrid fish (Fig 4A). However, while eye size was largely homogeneous in cave-surface hybrid fish that inherited a wild-type allele of *rx3* from their surface fish parent, cave-surface hybrid individuals that inherited a mutant allele showed a range of phenotypes, including having only one eye, having two eyes which were either both small or deformed, having two eyes with one resembling a wild-type eye while the other is reduced in size or deformed, or having no eyes at all (Fig 4A, Supplemental Fig 2).

**Figure 4.**
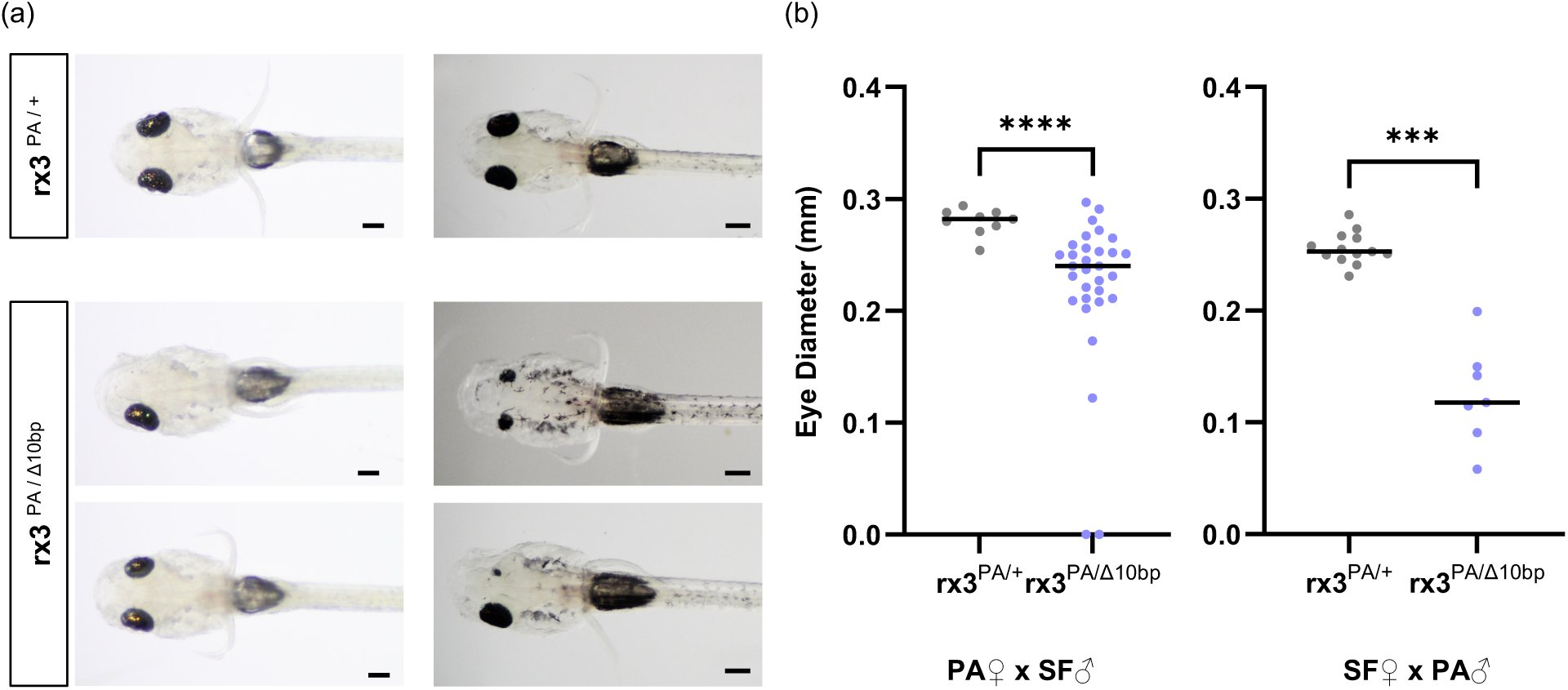
– Surface-cave hybrids exhibit a range in eye phenotypes. (a) Eye phenotypes observed in surface-cave hybrid fish at 5 days post fertilization. Fish are progeny from a cross between a Pachón cavefish and a surface fish that is heterozygous at the *rx3* locus for the engineered 10 bp deletion. Genotypes are indicated for each fish. Scale bars are 0.2 mm. Images on the left are from the cross from a Pachón female parent to a surface fish male and images on the right are from a Pachón male parent to a surface fish female. (b) Comparison of right eye diameter between hybrids that inherited a wildtype copy of the surface fish *rx3* allele (*rx3^PA/+^*) and hybrids that inherited a mutant copy of the surface fish *rx3* allele (*rx3^PA/Δ10bp^*). Mann-Whitney test: n=9,31; p-value= <0.0001 (Pachón female parent). Welch’s t-test: n=13, 7; t=7.504, p-value= 0.0002 (Pachón male parent). Individual data points indicate the eye diameter of individual fish and the lines indicate the median for the population. Crosses were made in two directions, with the Pachón fish as the female parent (left) or as the male parent (right), and offspring eye size was compared within crosses between siblings. *** = p<0.001, **** = p<0.0001.

Quantification of eye diameter in cave-surface hybrid fish revealed that eye size was quantitatively reduced in cave-surface hybrids that inherited the mutant surface fish *rx3* allele compared to their siblings that inherited the wild-type copy (Fig 4B, Supplemental Fig 3). These results were observed in reciprocal hybrids, both in progeny from crosses where the female parent was a cavefish and from crosses where the female parent was a surface fish (Fig 4B, Supplemental Figure 2, Supplemental Figure 3). These data support the hypothesis that cis-regulatory changes at the *rx3* locus contribute to eye size evolution in cavefish.

## 4. Discussion

### 4.1: Rx3 is required for eye development in A. mexicanus

The retinal homeobox (Rx/Rax) family of genes plays a critical role in early eye development. In mice, mutations in Rx/Rax prevent optic vesicle evagination, leading to eye loss (Mathers et al., 1997; Muranishi et al., 2012; Tucker et al., 2001). Moreover, in humans, mutations in Rax are associated with eye defects (Abouzeid et al., 2012; Lequeux et al., 2008; Voronina et al., 2004). In teleost fish, three *rx* genes have been identified (Chuang et al., 1999; Furutani-Seiki & Wittbrodt, 2004; Mathers et al., 1997), and mutations in the *rx3* gene in both zebrafish and medaka result in loss of the optic vesicles and eyes (Kennedy et al., 2004; Loosli et al., 2003, 2001; Winkler et al., 2000). Here, we investigate the consequences of mutations in *rx3*, a candidate gene for eye evolution in cavefish, in *A. mexicanus.* Similar to zebrafish and medaka, previous work demonstrated that *rx3* is expressed in the anterior neural plate in both surface and cave *A. mexicanus* (Agnès et al., 2022; Chuang et al., 1999; Sifuentes-Romero et al., 2020; Warren et al., 2021). In zebrafish, as development progresses, *rx3* expression is reduced in the eye primordia, and expressed in the developing hypothalamus, before being expressed in the retina as retinal differentiation occurs (Chuang et al., 1999). We found similar *rx3* expression patterns during eye development in *A. mexicanus* surface and cave fish. Further, we found that *rx3* is required for optic vesicle and eye formation in surface fish, as *rx3* homozygous mutant surface fish lack eyes, similar to the phenotypes observed in *rx3* mutant medaka and zebrafish (Kennedy et al., 2004; Loosli et al., 2003, 2001; Winkler et al., 2000). Together, these expression patterns and functional results in surface fish demonstrate that the role of *rx3* is conserved in *A. mexicanus,* and that *rx3* is required for eye development in this species.

### 4.2: Visually mediated behaviors are altered in rx3 mutant surface fish

Multiple behaviors that differ between cave and surface fish are altered when eyes are manipulated in surface fish, including larval prey capture behavior and schooling, implicating a requirement for vision for these behaviors in surface fish (Kowalko, Rohner, Rompani, et al., 2013; Lloyd et al., 2018). For example, while surface fish in the light capture prey head on, cavefish and surface fish in the dark capture prey by striking from the side, using their lateral line (Lloyd et al., 2018). However, while placing surface fish in the dark to assess behavior removes visual cues, fish can sense and respond to light in the absence of the eyes, for example, through the pineal gland. In cavefish, the pineal organ is intact and can function in mediating light-sensing behaviors (Yoshizawa & Jeffery, 2008). Thus, placing surface fish in the dark does not replicate the light-sensing state of cavefish. The establishment of a line of surface fish that lack eyes throughout development provides an opportunity to investigate the role of vision in behaviors that are altered between cavefish and surface fish in surface fish with a lack of vision due to genetic ablation of eyes. Further, as *rx3* mutant surface fish do not develop eyes, they provide the opportunity to determine if acute visual deprivation, such as placing fish in the dark, leads to the same behavioral shifts as raising fish in the absence of eye development.

We found that OMR is reduced in *rx3* mutant surface fish, similar to what was previously observed in larval cavefish (Choy et al., 2024). This suggests that the eyes are required for this behavior. Thus, defects in the eye itself in cavefish may be sufficient to result in loss of OMR. For example, while cavefish develop eyes, apoptosis in the lens begins shortly after optic cup formation (William R. Jeffery & Martasian, 1998). Further, while the retina develops in cavefish, there are defects in retinal cell types, including the photoreceptors, which lack outer segments (Emam et al., 2020). As loss of eyes through *rx3* mutation is sufficient to disrupt OMR in surface fish, these or other defects in the eye itself may contribute to loss of OMR in cavefish.

Similar to surface fish in the dark, eyeless *rx3* mutant surface fish have a significantly larger strike angle when hunting than their wild-type siblings, recapitulating the cavefish-like phenotype that is also observed in surface fish that have been acutely deprived of visual cues (Lloyd et al., 2018). This supports the hypothesis that altered striking behavior in cavefish occurs through the use of non-visually mediated cues in these animals. However, we did see other differences in prey capture between surface fish in the dark and *rx3* mutant surface fish. While previous work demonstrated that surface fish in the dark strike less than surface fish in the light (Lloyd et al., 2018), *rx3* mutant surface fish have a similar number of strike attempts compared to their wild type siblings. Whether this is due to *rx3* mutant fish learning to hunt without visual cues, or having enhancement of other sensory systems due to loss of eye development, remains to be investigated.

### 4.3: rx3 contributes to eye size evolution in cavefish via cis-regulatory changes

Multiple studies have used QTL mapping to identify loci associated with variation in eyes in *A. mexicanus.* Multiple QTL have been identified for a number of eye-related traits, including eye size, lens size, and thickness of the different retinal layers (Kowalko, Rohner, Linden, et al., 2013; Kowalko, Rohner, Rompani, et al., 2013; K. E. O’Quin et al., 2013; Protas et al., 2007, 2008; Yoshizawa et al., 2012). While mapping QTL to *A. mexicanus* genomes has revealed a number of candidate genes for eye evolution within these QTL (McGaugh et al., 2014; Warren et al., 2021; Wiese et al., 2024), causative genes and genetic variants within these QTL which underlie cavefish eye evolution remain largely unknown. An exception to this is the gene *cystathionine ß-synthase a* (*cbsa*). The *cbsa* gene was identified under an eye size QTL and expression of *cbsa* is downregulated in the developing cavefish eye (Ma et al., 2020). Loss of *cbsa* in surface fish results in eyes with ventrally displaced lenses or no eye formation at all due to defects in optic vasculature, suggesting that circulatory system changes due to the downregulation of this gene in the cavefish eye contribute to eye degeneration (Ma et al., 2020). Further, sequence analysis and evaluation of enhancer activity both suggest that cis-regulatory variation in *cbsa* underlies the role of this gene in cavefish eye evolution (Ma et al., 2020).

The *rx3* gene was identified as being located within QTL for the outer plexiform layer thickness of the retina and for eye size (McGaugh et al., 2014; K. E. O’Quin et al., 2013; Warren et al., 2021). Further, *rx3* expression is downregulated during eye field specification in the anterior neural plate in cavefish compared to surface fish (Agnès et al., 2022; Leclercq et al., 2024; McGaugh et al., 2014; Sifuentes-Romero et al., 2020; Warren et al., 2021). While sequence analysis suggests there are no protein coding changes at the *rx3* locus in cavefish (Agnès et al., 2022; McGaugh et al., 2014), analysis of allelic expression of *rx3* in F1 surface-cave hybrids suggests that cis-regulatory changes contribute to changes in *rx3* expression (Leclercq et al., 2024). However, the function of *rx3* in cavefish evolution was unknown. Here, we provide evidence that cis regulatory variation at the *rx3* locus contributes to eye size evolution in cavefish, as there is a lack of complementation in cave-surface hybrids that inherit the engineered 10 base pair allele from the surface fish parent.

How changes in *rx3* expression result in alterations to the eyes in cavefish is currently unknown. As expression of *rx3* after optic cup formation is qualitatively similar between surface and cavefish, and the expression domain size during eye specification is reduced in cavefish (Agnès et al., 2022; Sifuentes-Romero et al., 2020; Warren et al., 2021), one possibility is that cavefish have a smaller number of eye progenitor cells compared to surface fish. Future work identifying the cis-regulatory changes that contribute to these differences in *rx3* expression, as well as functional studies utilizing cave and surface fish *rx3* alleles, are required to determine the precise role of cis-regulatory variants in this gene in eye development.

## Acknowledgements

The authors thank all members of the Kowalko lab for help with fish care. This study was supported by National Institutes of Health grant R15HD009022 and R35GM138345 to Johanna Kowalkox.

