## Supplemental Figures for "The *rx3* gene contributes to the evolution of eye loss in the cavefish *Astyanax mexicanus*"

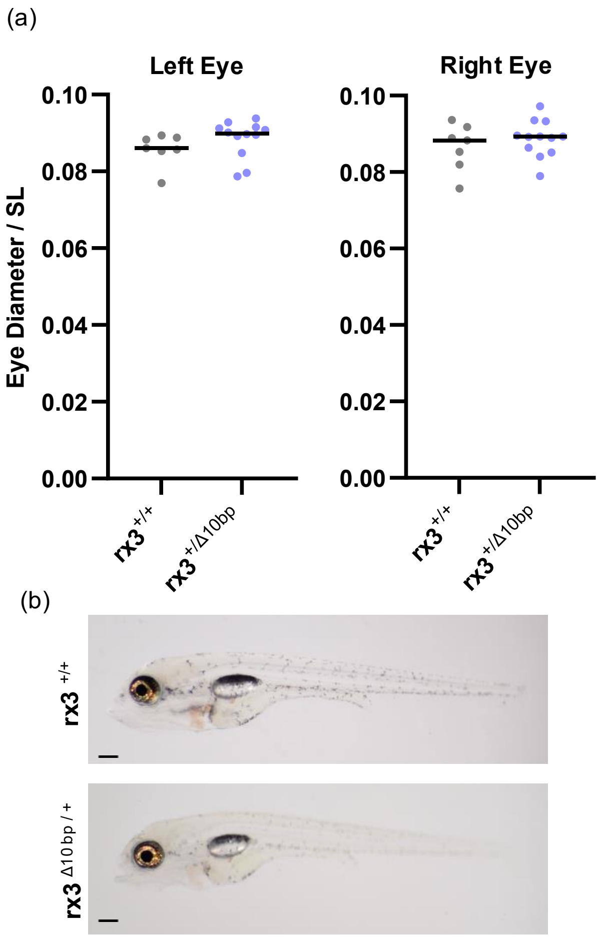


**Supplemental Figure 1- Surface fish that are heterozygous at the *rx3* locus have no significant differences in eye diameter compared to their wild-type siblings.** (a) Comparison of standardized eye diameter (eye diameter/standard length) between wild-type (*rx3*$^{+/+}$) surface fish and sibling surface fish that are heterozygous at the *rx3* locus (*rx3*$^{\Delta 10bp/+}$) Left eye: Mann-Whitney test: n=7,12; p-value= 0.0556. Right eye: Unpaired t-test: n= 7,12; p-value= 0.3885 (b) Images of representative *rx3*$^{+/+}$ and *rx3*$^{\Delta 10bp/+}$ fish. Scale bar is 0.2 mm.

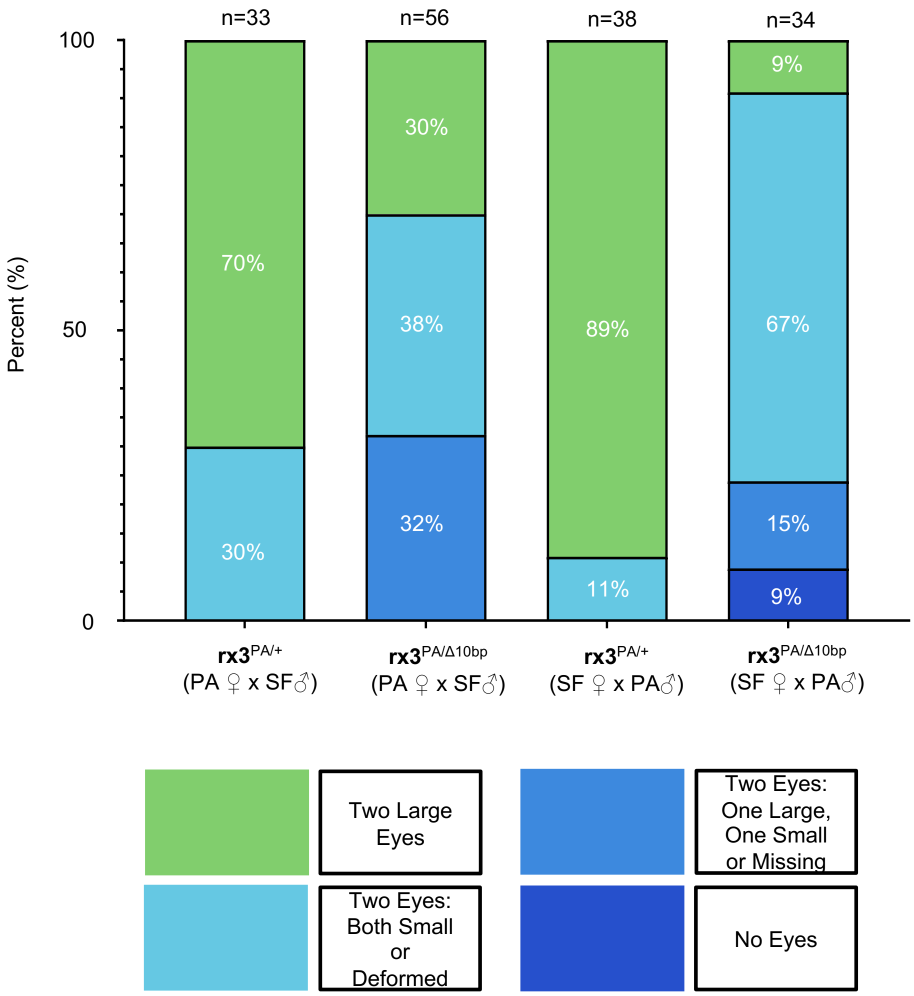


**Supplemental Figure 2 - Surface-cave hybrids that inherit a mutant copy of the surface fish *rx3* allele have a range in eye phenotypes**. Percentage of fish with an eye phenotype that falls within 4 possible categories – two large, two eyes that are both small or deformed, two eyes one of which is large and one of which is small or missing, or no eyes. Fish genotype as well as maternal identity (surface vs. Pachon cavefish) is indicated. + indicates inheritance of the wild-type allele from the surface fish parent. Δ10bp indicates inheritance of the mutant allele from the surface parent. PA indicates the Pachon cavefish *rx3* allele.

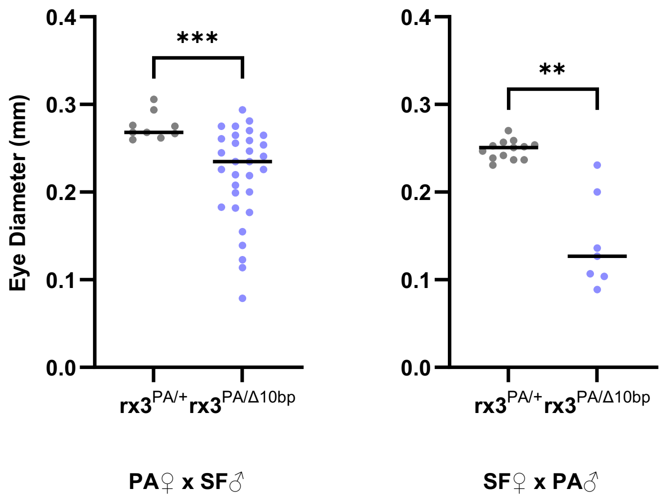


**Supplemental Figure 3 - Left size eye in cave-surface hybrid fish.**
Fish are progeny from a cross between a Pachón cavefish and a surface fish that is heterozygous at the *rx3* locus. Comparison of left eye diameter between hybrids that inherited a wildtype copy of the surface fish *rx3* allele ($rx3^{PA/+}$) and hybrids that inherited a mutant copy of the surface fish *rx3* allele ($rx3^{PA/\Delta 10bp}$). Mann-Whitney test: n=9,31; p-value= 0.0003 (Pachón female parent). Welch's t-test: n=13, 7; t=5.226, p-value= 0.0017 (Pachón male parent). Individual data points indicate the eye diameter of individual fish and the lines indicate the median for the population. Crosses were made in two directions, with the Pachón fish as the female parent (left) or as the male parent (right), and offspring eye size was compared within crosses between siblings.
